# A memetic algorithm enables efficient local and global all-atom protein-protein docking with backbone and sidechain flexibility

**DOI:** 10.1101/2021.04.12.437963

**Authors:** Daniel Varela, Vera Karlin, Ingemar André

## Abstract

Protein complex formation is encoded by specific interactions at the atomic scale, but the computational cost of modeling proteins at this level often requires the use of simplified energy models and limited conformational flexibility. In particular, the use of all-atom energy functions, backbone and sidechain flexibility results in rugged energy landscapes that are difficult to explore. In this study we develop a protein-protein docking algorithm, EvoDOCK, that combine the strength of a differential evolution algorithm for efficient exploration of the global search space with the benefits of a local optimization method to refine detailed atomic interactions. EvoDOCK enabled accurate and fast local and global protein-protein docking using an all-atom energy function with side-chain flexibility. Comparison with a standard method built on Monte Carlo optimization demonstrated improved accuracy and with increases in computational speed of up to 35 times. The evolutionary algorithm also enabled efficient atomistic docking with backbone flexibility.

## 1. Introduction

Interactions of proteins are critical for the function of most cellular processes (Alberts, 1998). While the number of experimentally determined protein complexes is steadily increasing in the Protein Data Bank (PDB) (Berman et al., 2000), it still covers only a sliver of the interactome. Prediction of the structure of protein complexes and assemblies, therefore, continues to be a central problem in structural biology (Lensink et al., 2017). Complex formation is the result of subtle energetic trade-offs in the bound and unbound states and is often associated with conformation changes in the binding partners. This results in rugged energy landscapes that are difficult to explore with protein-protein docking methods. In this study we introduce a new approach for conformational sampling at the atomistic level and show that it results in more efficient and accurate protein-protein docking simulations.

In many real-world applications of protein-protein docking, the starting point is the structure (models or experimental structures) of the individual interaction partners with an unknown binding mode. This is referred to as global docking (Huang, 2015). Identifying the correct binding mode involves a six-dimensional search of rigid-body orientations, where three degrees of freedom correspond to rotations and three encode the relative translation between the binding partners. The flexibility of sidechains and backbone can further expand the search space. To reduce the computational cost, protein-protein docking methods typically use a rapid, but less accurate, search strategy to reduce the number of docking orientations that are then evaluated and refined by more atomistic approaches using local protein-protein docking. Rigid body docking methods using Fast Fourier Transform (FFT) methods can be very fast. Furthermore, improvements of energy models that can be evaluated on a grid have enhanced their ability to identify good docking orientations (Desta et al., 2020b). An alternative strategy involves the use of coarse-grained models coupled to search strategies such as Metropolis Monte Carlo (MMC) (Gray et al., 2003) or evolutionary-based optimization methods (Torchala et al., 2020). The pool of docking orientations is then typically optimized with a refinement step, which couples the use of all-atom energy functions or force-fields with sidechain (and sometimes backbone) flexibility (Harmalkar and Gray, 2021).

The use of simplified energy representations has some benefits beyond computational speed. In particular, they are less sensitive to the approximation of a rigid backbone often used in protein-protein docking. On the other hand, their use can also lead to the exploration of docking orientations that are far from the native state. The true energy landscape is typically better represented with all-atom energy functions. However, skipping the initial global search step with FFT/coarse-graining methods and directly optimizing docking orientations based on an all-atom energy function have not been feasible due to the high computational cost. Nonetheless, we show here that such an approach can be feasible if a more efficient conformational search method is employed. By combining an evolutionary algorithm (EA) for global search and a local search method to refine solutions, we demonstrate how both accurate and relatively fast atomistic protein-protein docking can be carried out.

One of the main advantages of EAs is their ability to efficiently explore large areas of the global search space compared to a local search strategy like Monte Carlo. Local methods identify a new solution based on a small change of the current solution (a strategy known as “exploitation”). On the other hand, population-based search methods, such as evolutionary algorithms, maintain many feasible solutions during each search iteration or generation. This allows a global search in different areas of the space of solutions, with the possibility of “exploration” of new search areas through the genetic operators of the corresponding EA.

Both EAs and the local search can be combined in hybrid approaches known as “Memetic Algorithms” (MA), which aims to combine the exploration capability of an EA to identify the most promising regions of the search space while using local search to exploit the surroundings of the individual solutions (Moscato and Cotta, 2003). In recent years, MAs has been successfully applied to solve complex optimization problems in many scientific domains (Neri and Cotta, 2012) and been demonstrated to be a high-efficiency strategy to explore fitness landscapes ((Črepinšek et al., 2013; Lala et al., 2015)).

In this study, we present a method that uses the Differential Evolution (DE) algorithm as the evolutionary optimization framework and a MC-based docking approach as the local search method. DE (Storn and Price, 1997) (Price et al., 2005) is a population-based search algorithm that has been developed for optimization problems where possible solutions are encoded as real-valued vectors. This is the case in protein-protein docking where the relative position of two binding partners can be described by three translational and three rotational degrees of freedom. By converting DE into a memetic algorithm and combining it with a local optimization method, the search efficiency can be increased by refining the solutions proposed by the DE framework.

Several studies have presented evolutionary algorithms (such as genetic algorithms, swarm optimization and DE) to solve protein docking problems. Such approaches have been particularly employed for protein-ligand docking (Kang et al., 2009; de Magalhães et al., 2015; Narloch and Dorn, 2019). This includes some methods based on memetic algorithms, such as combinations of EAs with a variety of local search methods (Thomsen and Christensen, 2006; Chen et al., 2007; Grosdidier et al., 2009; Chung et al., 2013). In the context of protein-protein docking, evolutionary algorithms have been used for rigid-body docking (Gardiner et al., 2001; Sudha et al., 2019), while memetic approaches have also been defined (Leonhart et al., 2019; Torchala et al., 2020). Evolutionary algorithms can also enable exploration with back-bone diversity. For example, SwarmDOCK is a memetic algorithm based on swarm optimization, that includes conformational variability through normal mode analysis (Torchala et al., 2020).

These current applications of EAs to protein-protein docking have been limited to rigid-body sampling without sidechain and backbone flexibility and typically involve the use of simplified energy models and coarse-grained descriptions of proteins. The introduction of all-atom energy functions and conformational flexibility results in very rugged energy landscapes, which requires highly efficient search strategies to explore. In this study we present the protein-protein docking program EvoDOCK, which is based on a memetic algorithm where a differential evolution strategy is coupled to a Monte Carlo-based local search method built on RosettaDOCK (Gray et al., 2003; Chaudhury et al., 2011; Marze et al., 2018).

The results demonstrate that highly accurate local and global protein-protein docking predictions can be achieved by search directly using an all-atom energy function while simultaneously exploring sidechain flexibility. We also demonstrate that EvoDOCK can be extended to efficiently explore backbone flexibility during the atomistic docking.

## 2. Results

The EvoDOCK methodology for fixed backbone docking is schematically presented in Figure 1. An initial set of randomly generated docking orientations is prepared. This constitutes the initial population that is evolved over several generations. At each iteration of the Differential Evolution (DE) procedure, and for each current individual of the population, a trial solution is created by applying mutation and crossover operations on the parameters that describe the six docking degrees of freedom in the system based on the lowest energy individual. Each candidate is then optimized with a few cycles of a local refinement step in RosettaDOCK (Chaudhury et al., 2011), which involves a Monte-Carlo based optimization of the rigid-body orientation, a combination of greedy and combinatorial rotamer optimization and rigid-body minimization. After this local search process, the lowest energy solutions are selected to continue to the next iteration. After the final iteration, the model with the lowest Rosetta energy is selected as the result of an EvoDOCK run.

**Figure 1:**
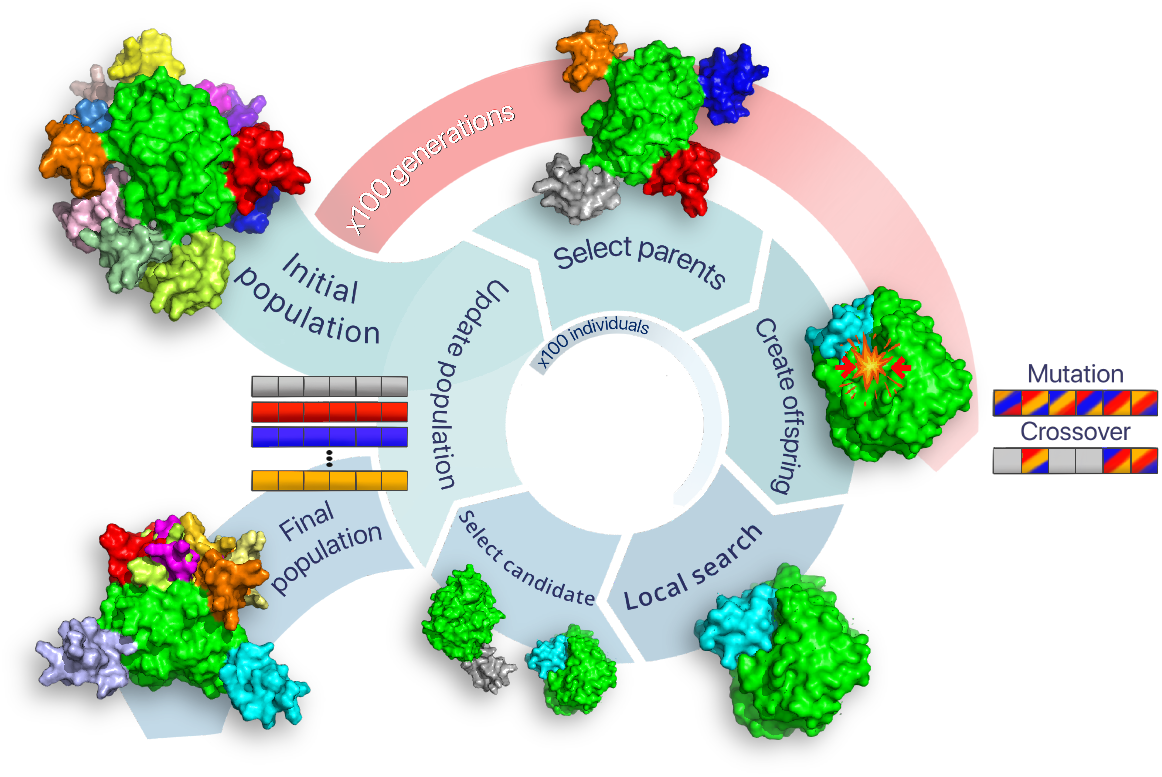
Schematic representation of the EvoDOCK protein-protein docking methodology. An initial population of 100 docking orientations is generated by randomly orienting binding partners around the green static protein. Out of the population, lowest energy individual and two parents are randomly selected to create a new offspring, illustrated here by the orange, blue and red members. The parents are combined to form the cyan offspring using mutation or crossover operations for each of the six docking degrees of freedom. This candidate typically has clashes that need to be resolved by a local search involving RosettaDock, which optimizes the rigid-body orientation and sidechain conformations. Two Monte Carlo cycles of RosettaDOCK is called in the default EvoDOCK protocol. The energy of the candidate is then compared to one of the selected parents (grey in this example) to determine whether it will replace it in the population. Each member of the population is compared with a candidate offspring in each generation. After 100 generations the lowest energy model is selected as the final prediction.

In the sections below we characterize EvoDOCK in global and local docking experiments, compare it with an all-atom and a FFT-based docking program, and evaluate docking efficiency. Finally, we extend EvoDOCK to handle flexible backbone docking and apply it to a challenging modeling scenario where binding induces considerably backbone adjustments.

### 2.1. Global docking performance of EvoDOCK

The performance of EvoDOCK was tested in global docking experiments in which the native binding mode is recovered without prior knowledge of the binding site and sidechain conformations in the binding interface. Since the focus in this section is to evaluate improvements in general search efficiency, we created a benchmark with complexes where the Rosetta all-atom energy function is able to identify the native as the energetically most favorable binding mode and where backbone adjustments are not required to identify the optimal solution (bound-bound docking). To compare the performance of EvoDOCK we also carried out global docking calculations using the standard rigid backbone docking protocol in RosettaDOCK. RosettaDOCK combines a Monte Carlo-based rigid body optimization in a coarse-grained representation followed by all-atom Monte Carlo optimization. The stochastic nature of the docking calculations requires collection of a large number of docking trajectories to get sufficient statistical basis for our conclusions and this comes with a substantial computational cost. Hence, we limit ourselves to 10 protein com-plexes and study them in detail. The complexes were selected based on the results in the RosettaDOCK local docking benchmarking study (Chaudhury et al., 2011) and are shown in Figure S1.

The solution space accessible to EvoDOCK was comprehensively mapped by running 1,000 independent docking runs. This is considerably more than what is typically required to identify the native binding mode but enables the collection of appropriate statistics on docking performance. In total, a number of 100,000 (100×1000) members were evolved in these docking calculations for each complex. The same number of models was generated by Rosetta-DOCK for comparison with each complex. The docking energy landscape was summarized as the scatter plots in Figure 2, where the energy of each model was plotted against the interface RMSD (iRMSD) to the native complex. In all of the 10 cases, the 1,000 models predicted by EvoDOCK have low energy values and a significant fraction of models have low iRMSD values. Figure 2 also presents the complete evolved population for each benchmark complex (100 from each independent run). The range of iRMSD values sampled in this population are broader, but all members still have low energy values. In contrast, RosettaDOCK solutions have a wide range of energy and iRMSD values. High energy solutions are common and only a small fraction of models are typically sampled in the near-native area. For example, RosettaDOCK generates docking solutions with iRMSD values up to 500Å for the *1b6c* complex, and less than 2% of solutions have iRMSD values lower than 10 Å. The difference between the lowest and the highest energy values has a spread of more than 1,000 energy units. In contrast, EvoDOCK solutions converge to a narrow distribution near the native conformation. The maximum iRMSD value among the 100,000 models produced by EvoDOCK was 12 Å, with 94 % solutions having iRMSD values lower than 10 Å and the largest spread in energy less than 27 energy units. Similar results were observed for all the benchmark cases.

**Figure 2:**
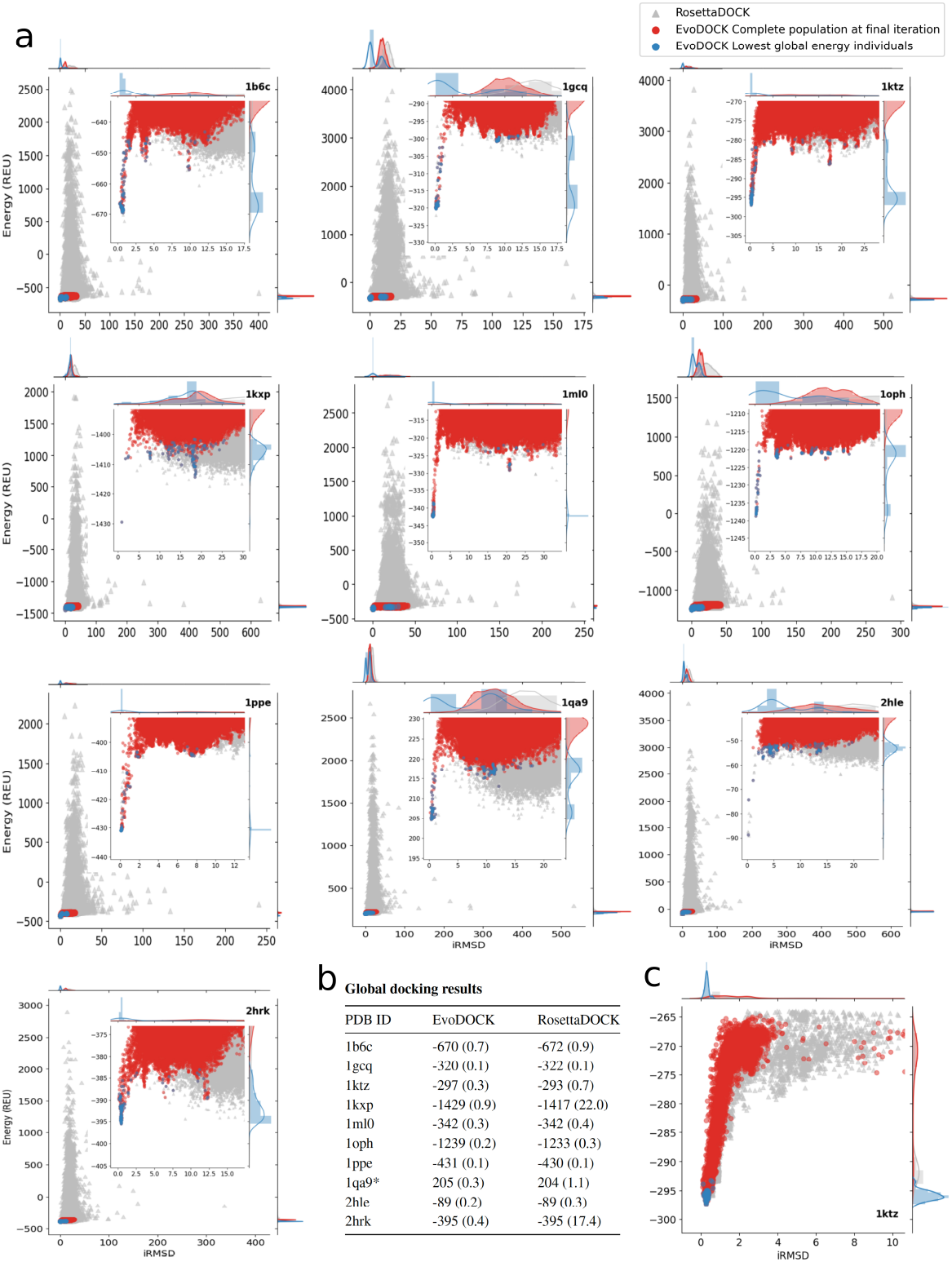
Distribution of the global docking results for 100,000 models. **a**: Global docking results with EvoDOCK and RosettaDOCK. Each model is represented by a point where Rosetta full-atom Energy Units (REU) are shown on the *y*-axis with the corresponding interface RMSD (iRMSD) on the *x*-axis. The blue points are the result of 1,000 independent evolutionary processes by EvoDOCK, while the red points are the complete EvoDOCK population at the end of the trajectory (100,000 models). The RosettaDOCK models resulting from 100,000 independent trajectories are shown in grey. The inset shows a zoom into the region of low iRMSD to visualize the details of “near-native” decoys. **b**: Comparison of the lowest energy value obtained for each benchmark complex and its corresponding iRMSD (in parentheses) for EvoDOCK and RosettaDOCK. **c**: Exploration of the energy landscape with the local docking for the benchmark complex *1ktz*. The 5,000 models obtained in the optimized population by EvoDOCK are shown in red (the lowest energy individual achieved by each of 50 runs is represented in blue) and the 5,000 RosettaDOCK models are shown in grey for comparison. **1qa9\***: Complex *1qa9* shows a high energy value as a consequence of a backbone clash that is in the core of one of the binding partners.

Focusing on a region near the native binding mode, we observe that EvoDOCK produces more models in this range than RosettaDOCK. In particular, below 2 Å iRMSD, EvoDOCK samples more models than RosettaDOCK in 9 out of 10 cases (Table S1). With the exception of two complexes, 4 to 22% of all models predicted by EvoDOCK end up in this range, while RosettaDOCK never goes above 0.3%. In terms of the lowest energy identified in the simulations, the results are more similar. In 4 cases (*1ktz*, *1kxp*, *2hrk* and *1oph*) EvoDOCK produces the lowest energy model, RosettaDOCK finds the lowest energy in 3 cases (*1b6c*, *1gcq* and *1qa9*), while for the remaining 3 cases the energies are comparable.

In a blind docking experiment where the native structure is not known, solutions would be ranked by energy. Therefore, the quality of the docking results can also be assessed by comparing the iRMSD of the lowest energy solution. The table in Figure 2b summarizes these results. In all the cases, the lowest energy model obtained by EvoDOCK has a lower or equal iRMSD than RosettaDOCK. For two complexes, *1kxp* and *2hrk*, RosettaDOCK is not able to predict the native binding mode. In the one case where RosettaDOCK produces a better prediction (*1gcq*), the difference to EvoDOCK is insignificant (0.03 Å).

### 2.2. Increase in computational efficiency of EvoDOCK

The results presented so far demonstrate that EvoDOCK performs a detailed sampling near the native binding mode. But an important characteristic of a docking method is also the computational cost. In a Monte Carlo-based docking algorithm, each run can be relatively short but it is necessary to sample a large number of different independent trajectories in order to sample near-native solutions. In contrast, an evolutionary algorithm optimizes several trajectories simultaneously and takes longer to complete, but with potentially fewer runs. To determine the number of EvoDOCK runs and aggregated simulation time necessary to achieve high-quality docking results, we have to define a metric for success. We considered a successful docking experiment one in which the lowest energy prediction has an iRMSD of 2.0 Å or less to the native binding mode, in which the molecular interactions are likely to be very similar to the native complex. Since docking simulations are stochastic, we need to define a probabilistic measure. *P_success_* measures the probability of success (iRMSD ≤2.0 Å for lowest energy prediction) with a given number of models generated. To evaluate *P_success_*, we sampled data sets using a bootstrap method based on the 1,000 EvoDOCK runs (with a total of 100,000 models) for each complex, following the strategy described by Chaudhury et al. (Chaudhury et al., 2011).

In Figure 3, *P_success_* is plotted against computational time (in hours) on a single core of an Intel Xeon E5-2650 v3 processor (2.3 GHz). For comparison we have evaluated the same metric on the RosettaDOCK simulations. As it can be seen in Table S2, in 9 out of 10 cases, given the same *P_success_*, EvoDOCK reduces the necessary computational time in comparison to Roset-taDOCK. The docking results can be divided into three categories. In the first category, both EvoDOCK and RosettaDOCK successfully produce high-quality docking solutions and with *P_success_* values up to 100% (*1b6c*, *1gcq*, *1ktz*, *1ml0*, *1ppe*). For these cases, we can compare computational efficiency by comparing the computational time required to achieve a specific probability of success. In all these cases, EvoDOCK provides a significant speed boost from 4 to 38 times faster. A second category corresponds to the cases where EvoDOCK succeed in reaching *P_success_* values, while RosettaDOCK plateaus at lower values. For example, for *1qa9*, RosettaDOCK achieves a maximum *P_success_* of 65%, a level that EvoDOCK can reach more than 30 times faster. Similarly, the maximum *P_success_* for complex *1oph* for RosettaDOCK is near 90%, which can be reached 15 times faster by EvoDOCK. In the third category of results, RosettaDOCK cannot identify any near-native solutions (*2hrk* and *1kxp*). This is probably due to enhanced sampling around false minima at high iRMSD, which can be seen in Figure 2. EvoDOCK finds near-native solutions, but achieves only a *P_success_* of 65% with *1kxp*, suggesting that this complex has a difficult landscape to explore. Finally, for one case, *2hle*, Roset-taDOCK is more efficient at the sampling binding mode.

**Figure 3:**
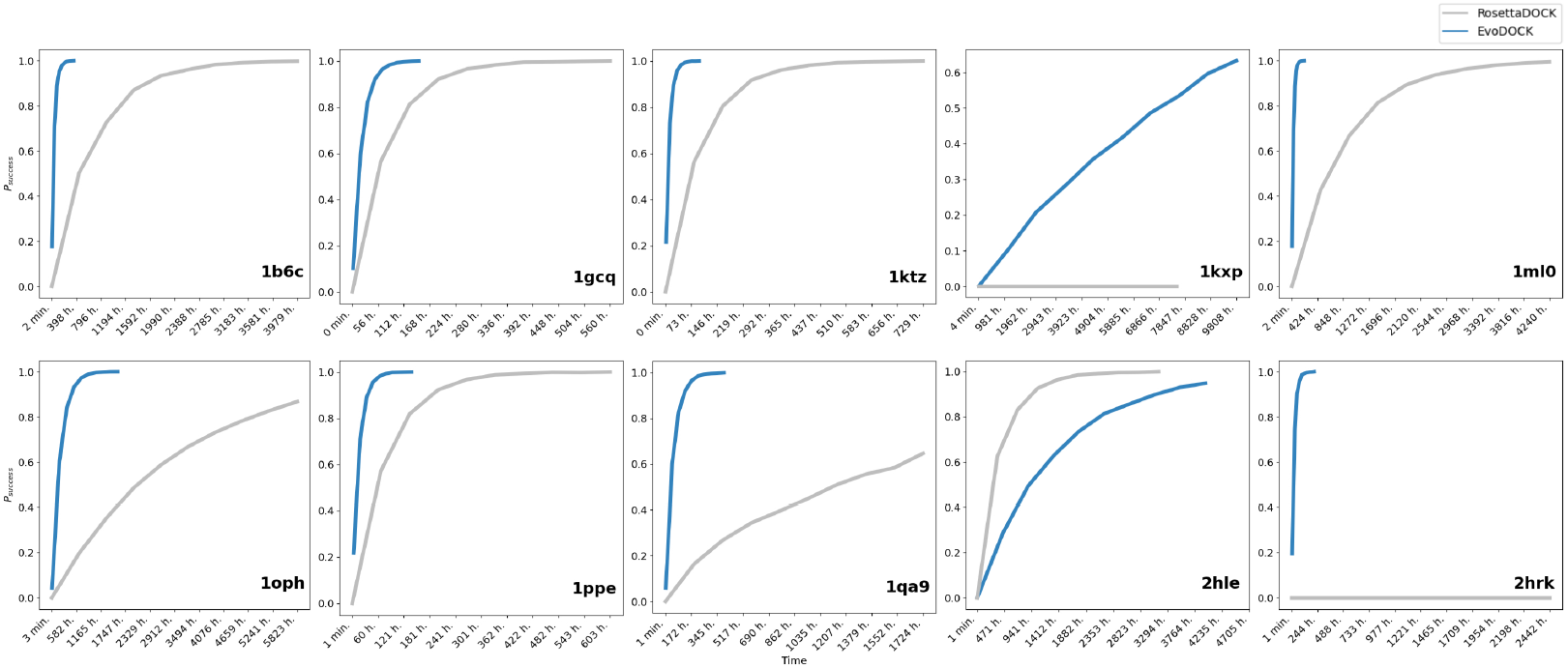
Computational efficiency comparison of EvoDOCK and RosettaDOCK. Comparison of the efficiency of EvoDOCK and RosettaDOCK in identifying “successful” docking orientations. A prediction is defined as “successful” when the iRMSD of the lowest energy individual is below a threshold of 2.0 Å. Plots show the value of *P_success_* (*y*-axis), obtained by bootstrapping, for sampled sets of predictions as a function of computational time (*x*-axis). Computational time was calculated by taking the average time for completing one run/model and multiplying it with the total number of runs/trajectories for EvoDock/RosettaDOCK respectively.

We can also quantify the number of runs needed by either method to achieve a *P_success_* value higher than 80%, see table 1. EvoDOCK needs an average of 18 runs in order to reach the *P_success_* threshold value for 8 out of 9 cases this threshold is reached, with 100 solutions (population size) minimized simultaneously during each run. In comparison, RosettaDOCK requires an average of 29,500 independent Monte Carlo trajectories to reach the same threshold.

**Table 1:**
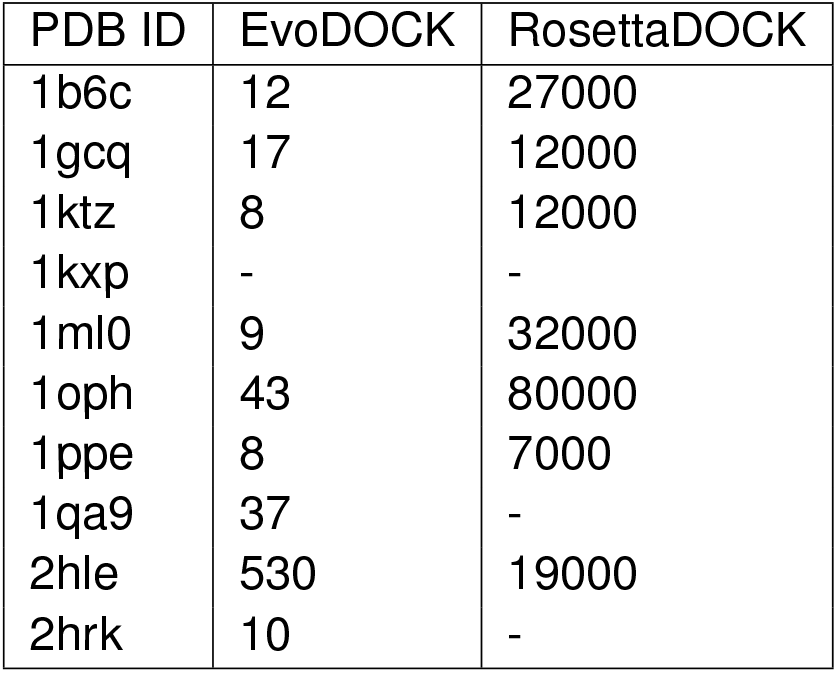
Number of independent runs necessary to achieve a *P_success_* of 80% with EvoDOCK and RosettaDOCK. Each EvoDOCK run involves a population of 100 models.

| PDB ID | EvoDOCK | RosettaDOCK |
| --- | --- | --- |
| 1b6c | 12 | 27000 |
| 1gcq | 17 | 12000 |
| 1ktz | 8 | 12000 |
| 1kxp | - | - |
| 1ml0 | 9 | 32000 |
| 1oph | 43 | 80000 |
| 1ppe | 8 | 7000 |
| 1qa9 | 37 | - |
| 2hle | 530 | 19000 |
| 2hrk | 10 | - |

### 2.3. Comparison with FFT-based global docking

Global protein-protein docking is typically carried out by FFT-based methods like ZDOCK (Chen et al., 2003), GRAMM (Tovchigrechko and Vakser, 2005), pyDOCK (Cheng et al., 2007) and ClusPro (Kozakov et al., 2017). We selected ClusPro to compare with EvoDOCK since its one of the most successful automatic methods in CAPRI over the last decade and is one of the most popular docking servers (Desta et al., 2020a). The FFT calculation in ClusPro is carried out by PIPER (Kozakov et al., 2006). Low energy models are then clustered and the center of most populous clusters are then energy minimized using the CHARMM potential (Brooks et al., 1983). In the Table S7, the iRMSD of cluster center of the most populous cluster is presented using the ‘Balanced’ scoring scheme during the FFT. The values range is between 1.3 and 23.1 Å, with 6 out of 10 predictions having over 2 Å iRMSD. Results are similar with the other scoring schemes applied with the program. However, we can also look at iRMSD of all the models in all the populous clusters. The minimum iRMSD values vary between 1.1 and 2.4 Å in this set. This demonstrates that ClusPro samples low iRMSD binding modes, although the challenge is to pick them out based on the energy and clustering metrics used to rank models.

We then compared the computational time with the FFT-based ClusPro. Since the timings were not available from the ClusPro server we ran docking calculations with FFT program PIPER, which is the most time-consuming part of ClusPro. The results are found in table S7 and show that PIPER needs an average computational time for the four different scoring schemes used by ClusPro of 35 hours. On the other hand, EvoDOCK, to achieve a *P_success_* of 60% and 80%, requires a total computational time of 45 and 78 hours, respectively. That is a significant improvement compared to the more than 800/1000 hours of computational time that RosettaDOCK needs to achieve a *P_success_* of 60/80%, respectively.

One interesting application of EvoDOCK is to refine global docking solutions provided by FFT-docking. This combines the strengths of speed and coverage of FFT global docking, with the efficient optimization of atomistic interactions with EvoDOCK. To evaluate this approach we selected the complex that was most challenging for ClusPro (*1ml0*) and used its 92 number of solutions as starting rigid body positions to a EvoDOCK run. We found that EvoDOCK local search (running 10 independent simulations) is able to minimize the energy values and achieve a minimum iRMSD of 0.3 Å, while the best input ClusPro model had an iRMSD of 10.0 Å.

The complexes were selected on the criterion that the Rosetta energy function can identify the native conformation with lowest energy value, therefore not giving a level playing field with the ClusPro method. In order to evaluate the comparison in a unbound-unbound docking scenario, we selected a random set of 10 complexes with different difficulty categories in the Docking Benchmark v5.5 presented by the Weng group (Vreven et al., 2015) and evaluated the modeling results with DockQ (Basu and Wallner, 2016), a quality criteria commonly used at CAPRI experiments. Results are summarized at Table S8. Looking at the 100 lowest energy models of EvoDOCK and for all cluster centers and all four scoring schemes for ClusPro, the results are similar for both methods. For 3 (EvoDOCK) vs 4 (RosettaDOCK) complexes, an iRMSD below 4 Å is sampled, so few high quality predictions are made in this set. However, one of the EvoDOCK predictions obtains a DockQ score of 0.87 and iRMSD of 0.45 Å, and this is also the lowest energy prediction of the sampled models. The corresponding value for ClusPro among all the cluster centers is a DockQ score of 0.69 and iRMSD of 1.63 Å, with the cluster center selected by balanced score scheme having a DockQ score of 0.03 and iRMSD of 11.83 Å. This highlights that in order for unbound-unbound rigid-body docking to be successful, there must be compatible backbones. This is rarely the case, suggesting the need for a flexible backbone docking approach (see section 2.5).

### 2.4. Local Docking results

Local docking refinement serves as an important complement to global docking. It enables the improvement of solutions discovered in global docking simulations, the refinement of coarse-grained docking solutions, the characterization of the local energy landscape and the discovery of energy funnels. To evaluate EvoDOCK in the context of local docking refinement, the method was adapted to enable sampling around a given starting point, rather than from random positions as in the global docking scenario. In the local docking simulations, the initial positions are randomly perturbed around a given starting point by adding randomly selected rotation and translation perturbations. After this initiation, the methodology is the same as in the global docking case. Experiments were carried out using the same benchmark complexes used in the global docking, starting from the native binding mode and RosettaDOCK simulations were used as reference simulations using the same set of maximum perturbations. For EvoDOCK, we filtered out the starting states with less than 2Å to the native binding mode so that the initial population did not include the desired binding mode.

For both EvoDOCK and RosettaDOCK, local refinement results converge to near-native binding modes (Figure S2). As can be observed in table Table S3, both methods achieve a very similar result in terms of energies and iRMSD values, with dense sampling around the native binding mode. To map out the energy landscape around the native binding mode more comprehensively, the complete population obtained at the last iteration of the evolutionary process can be analyzed. This is demonstrated in Figure 2c for the complex *1ktz*. The additional models highlight alternative minima in the docking landscape not found in the energy-optimized EvoDOCK solutions. When all local refinement results are evaluated with this approach (Figure S2) we observe that EvoDOCK is able to densely sample the local docking energy landscape.

### 2.5. Flexible backbone Docking

**Figure 4:**
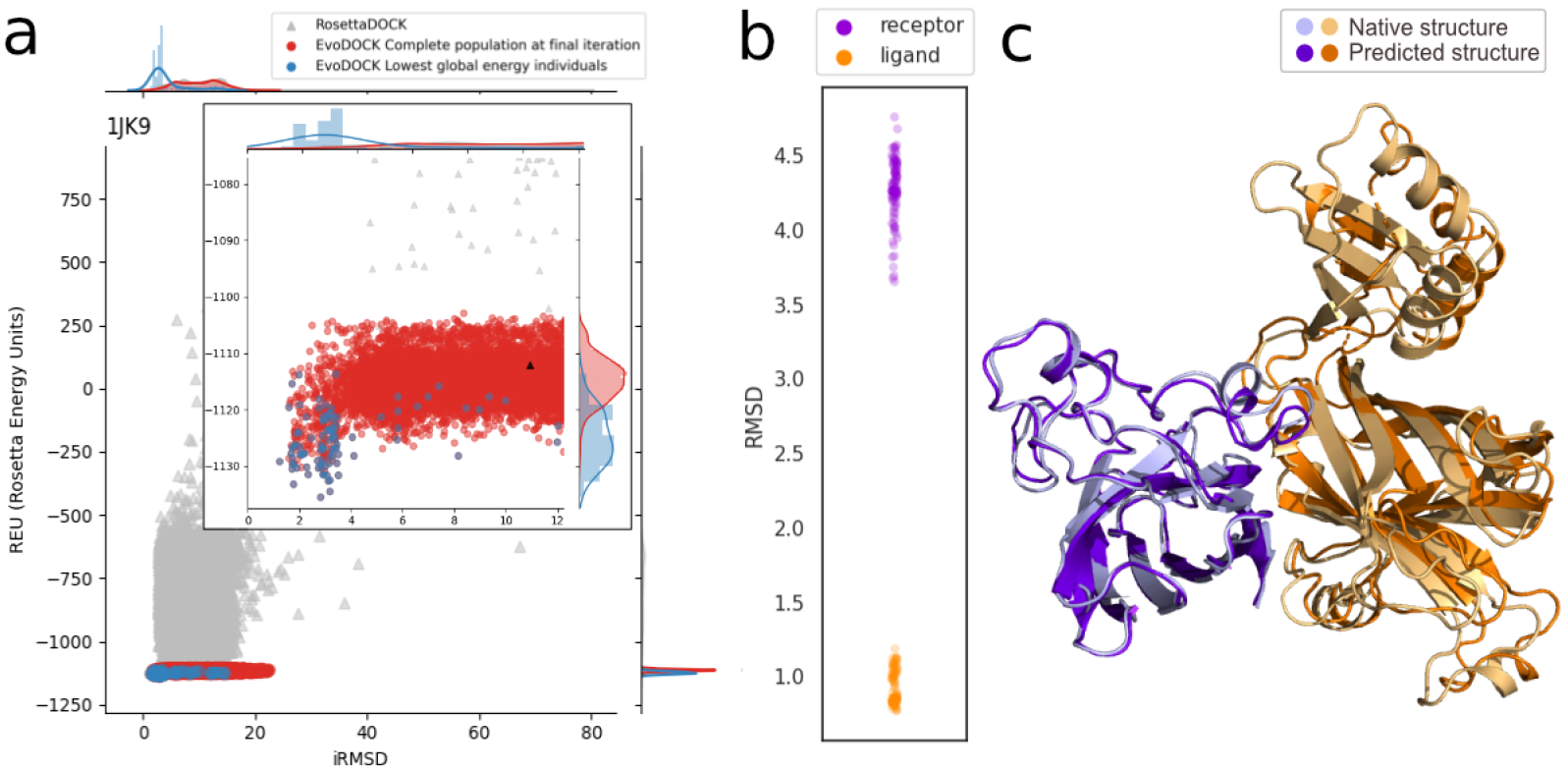
Exploration of the energy landscape with unbound flexible docking for the complex superoxide dismutase and its metallochaperone (pdbid *1jk9* for the bound complex, *2jcw* and *1qup*, for the unbound receptor and ligand, respectively) (Figure S1). Panel *a* contains a plot of Rosetta energy (*y*-axis) versus interface RMSD (iRMSD, *x*-axis) for each sampled model. The 10,000 models obtained in the optimized population by EvoDOCK are shown in red (the lowest energy individual achieved by each of 100 runs is represented in blue) and the 10,000 RosettaDOCK models are shown in grey for comparison (lowest energy individual highlighted in black). In panel *b*, the RMSD for each of the 100 generated ensembles to the bound conformation is shown (receptor in purple and ligand in orange). Panel *c* show the lowest energy individual obtained with EvoDOCK (dark) and native complex (light).

Complex formation is often associated with backbone adjustments in the binding partners upon association. Docking strategies based on rigid body approximations often struggle with such systems, in particular if an atomistic energy function is utilized. The evolutionary algorithm in EvoDOCK can readily be adjusted to allow for backbone flexibility using three different strategies: The local search can be complemented by an all-atom refinement step, a library of backbone conformations can be utilized or these two strategies can be combined. The conformational library approach is based on the method used by RosettaDOCK (Marze et al., 2018). RosettaDOCK uses a pregenerated backbone library of 100 conformers generated by all-atom refinement, back-rub sampling and normal mode analysis. During the coarse-grained docking conformers are sampled and the probability of conformers swaps is dynamically adjusted to reduce computational cost.

We implemented a method for backbone sampling in EvoDOCK where backbones are randomly selected and swapped with some probability at the end of a generation in the evolutionary algorithm. To evaluate the performance of EvoDOCK in sampling of backbone flexibility we selected a set of ten complex categorized as difficult in the Docking Benchmark v5.5 (Vreven et al., 2015).

In Figure S5, the 10,000 flexible backbone models are shown for EvoDOCK and RosettaDOCK. Similarly to what was observed for fixed backbone, EvoDOCK produces models with substantially better energies within a more narrow range of lower energy models and iRMSD values. EvoDOCK solutions have lower energy and improved DockQ score in 8 out of 10 cases (Table S9), while the iRMSD of the lowest energy is lowest for 9 of the cases. EvoDOCK results show a funnel-like shape energy landscape in the complexes *1eer*, *1fq1*, *1rke*, *1r8s* and *1jk9*. In Figure 2.5*a* we show one example, the complex between superoxide dismutase and its metallochaperone, where EvoDOCK shows a funnel towards a near-native state, whereas RosettaDOCK cannot sample high quality models with reasonable energies. The lowest energy model of EvoDOCK has a iRMSD of 2.8 Å while the best energy prediction for Rosetta-DOCK has an iRMSD of 10.9 Å. DockQ score presents a difference of 0.27 (0.81 and 0.54 for EvoDOCK and RosettaDOCK, respectively). However, both methods fail to identify a very accurate models of the complexes (1 out of 10 cases with high quality CAPRI category for EvoDOCK and zero for Rosetta-DOCK). This is expected due to the structural differences of the ensembles with the bound conformation (Figure S6). As it is shown in Figure 2.5*b* for the superoxide dismutase complex, none of the 100 conformers in the ensemble of the receptor has a RMSD below 3.6 Å relative to the bound conformation.

In summary, our result demonstrates that the increased accuracy of EvoDOCK in conformational sampling can also be observed when backbone flexibility is utilized.

## 3. Discussion

This study presents an alternative sampling strategy to improve the efficiency of conformational search for the computationally challenging problem of protein-protein docking. In contrast to the local search strategy provided by independent runs of MMC, DE is a population-based search method, which allows for information exchange between different solutions. As indicated by Feoktistov (Feoktistov, 2006), the fundamental idea of the algorithm is that it intrinsically balances exploration and exploitation in the search by modifying solutions by vector differences between current solutions. At the early iterations, the exploration is large because individuals are very different from each other. As the evolution proceeds, the population is guided by the funnel-shape of the Rosetta energy landscape and converges towards a minimum, and the smaller differences between individuals lead to a stronger emphasis on exploitation which results in refinement of solutions.

The use of standard mutation and recombination operators, however, can lead to unfeasible candidate solutions with clashes in the backbone or sidechains. To address this issue, EvoDOCK uses a memetic strategy (Moscato and Cotta, 2003), where DE solutions are repaired by a local refinement method that resolves backbone clashes and optimizes sidechain conformations. As detailed in Figure S3, this hybrid approach is essential for providing the evolutionary selection process with low energy candidates and in order to progressively optimize the solutions of the population. The default choice of two refinement steps per model provides a good balance between computational speed and accuracy. With a single refinement step, the prediction accuracy is somewhat reduced but is still comparable (Table S4). This is associated with an average 20% reduction in computational time, but the decrease is not uniform across all the benchmark complexes, with some complexes taking longer time with one rather than two refinement steps (Table S5). This suggests that the local refinement step also contributes to the convergence of the evolutionary algorithm. In the absence of the local refinement stage, the DE algorithm plateaus at high energy values (Figure S4). Increasing the number of DE iterations in order to obtain the same computational cost as with runs using the local refinement step cannot compensate (Figure S5). Hence the combination of DE and local refinement by RosettaDOCK is critical for the performance of EvoDOCK.

Our study on flexible backbone docking illustrate that the evolutionary algorithm in EvoDOCK can be used to efficient exploration of the energy landscape and with only minor additional computational costs. Collectively, our results show that EvoDOCK is very good at picking up the optimal solution if compatible backbones are available. However, to achieve better accuracy in prediction of flexible backbone targets, increased sampling efficiency is only part of the answer. Improved methodology for generating initial conformational diversity and refinement of backbones of protein complexes is equally important, something that is not explored in this study.

The recent revolution in the accuracy in protein structure prediction with deep learning methods (Jumper et al., 2021; Baek et al., 2021), will have significant implications for protein-protein docking by providing more accurate starting models. Nonetheless, a protein-protein docking method like EvoDOCK is still required for complex prediction. Methods like AlphaFold-Multimer (Evans et al., 2021) can be used to directly predict complexes, but refinement of the docking position is likely to be neccesary for accurate predictions in many cases. In summary, we demonstrate that a memetic docking algorithm can explore the rugged energy landscape of protein docking highly efficiently in the presence of both sidechain and backbone flexibility. The relative simplicity and flexibility of the methodology makes it an appropriate base for the development of more complex docking protocols with a possibility of custom tailored protocols towards specific systems of interest.

## Acknowledgments

The computations were enabled by resources provided by the Swedish National Infrastructure for Computing (SNIC) at HPC2N partially funded by the Swedish Research Council through grant agreement no. 2020/5-308 and the compute center LUNARC at Lund University with agreement no. 2021/2-59.

This work was supported by the European Research Council (ERC) under the European Union’s Horizon 2020 research and innovation program [Grant agreement No. 771820].

## Author Contributions

D.V. developed the code. D.V. and V.K. simulations and data analysis. D.V. and I.A. analyzed the data and wrote the manuscript.

## 4. Methods

### 4.1. Preparation of structures

The input complexes were prepared by the docking prepack protocol in Rosetta (Chaudhury et al., 2011) to remove clashes between the sidechains in the experimental structure, using a command line found in the Supplementary Information. The sidechain repacking is done when the binding partners are separated so that the sidechain conformations in the binding interface are not kept in the input structure for docking. This operation modify the structure in a RMSD range of [0.2, 1.1] Å.

### 4.2. Positioning of binding partners in EvoDOCK

In protein-protein docking, one protein partner is normally fixed (known as the “static molecule”) and the other protein partner moves in relation to the static protein (the so-called “moving molecule”). The relative orientation between the binding partners can be specified by a six-dimensional vector, where three values correspond to a rotation (*ψ*, *θ*, *φ*) and three to a translation (*x*, *y*, *z*). The rotation is specified by three Euler angles obtained by the orientation of the “moving molecule” with respect to a fixed coordinate system. The translation is defined in relation to the relative position of the “static molecule”. Rotation values are sampled at a maximum range of [-180,180], while translation values during the evolutionary process are sampled in the following way:

1. The moving molecule is randomly placed with respect to the rigid molecule at a large “non-contact” distance.
2. The local search method calls a simple procedure to slide the moving molecule into atomic contact with the static molecule.
3. The ensemble is analyzed to identify the largest value of *x*, *y*, and *z* in the translation vector after the local search. These are used as the upper bound in the evolutionary process for the translation vector.

This procedure ensures that the initial solutions are close enough to the “static molecule” and that each *x*, *y* and *z* translation is constrained by a reasonable value.

### 4.3. The Memetic Differential Evolution

The algorithm begins by creating a random set of candidate solutions defined as *N*umber of *P*arents (*NP*). The six-dimensional vector containing the information about rotation and translation of each candidate is randomly sampled at the ranges described in the previous section. That is, the genotype of each solution corresponds to the six dimensional vector. Additionally, these values are normalized in the interval [-1,1] for use with the DE genetic operators, which is frequently used in classic implementations of EAs. When the fitness is assigned, the encoded parameters are decoded to their corresponding values.

The standard implementation of DE algorithm (Price et al., 2005) is employed to optimize the solutions guided by a fitness function. In our docking calculations, the fitness function corresponds to the Rosetta full-atom energy function (*ref2015*). In order to create new candidate solutions, DE uses the mutation and crossover operators following the procedure defined in Algorithm 1. The method starts by selecting three different vectors (randomly chosen) from the current population. Then, for each of the 6 degrees of freedom (DoFs), given a crossover probability(*CR*), the candidate parameters are selected over a given vector of the population (target vector *P*[*i*]) and a “mutant vector”. The mutation operation creates a difference from vectors (*P*[*i*_2_][*j*] – *P*[*i*_3_][*j*]) multiplied by factor (*F*) and then added to the base vector *P*[*i*_1_]. The differential evolution strategy configuration used is *DE*/*bin*/*1*/*best*, where “*bin*” indicates that binomial crossover is employed during the formation of the trial population and “*best*” that the lowest energy individual is used as the base vector for the mutation operation, while the remaining two vectors are selected randomly. The aforementioned *F* and *CR* parameters are used to control the search performance (the exploration and exploitation). During our experiments, we have used a fixed set configuration where *F* = 0.9 (usually between [0,2]) and *CR* = 0.3 (usually in range [0,1]), determined experimentally to obtain a balanced search between exploration and exploitation.

#### Algorithm 1 Create candidate O[i] from parent P[i]

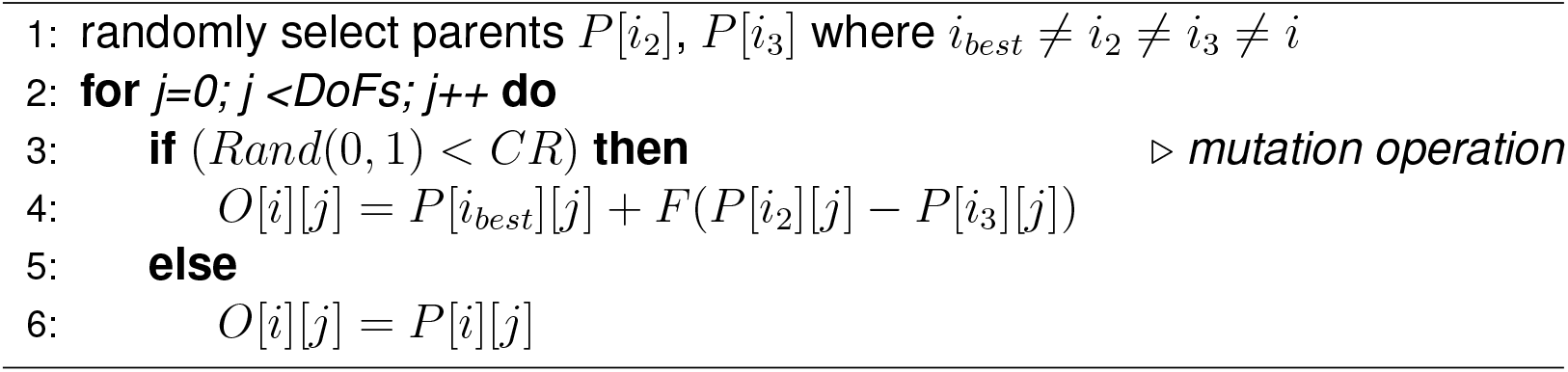

In the memetic algorithm, DE is combined with a local search (LS) applied to every candidate, where the local search operator acts as a repair mechanism and a refinement method for the candidate. Further details of the effect of LS during the evolutionary process are shown in the Supplementary Information, Figure S3. The local search operator starts with a *slide_into_contact* method provided by Rosetta, where the “moving molecule” is translated along a vector at the center of mass until it contacts the static body, followed by two cycles of a rigid-body MMC energy minimization and a combinatorial sidechain optimization. As in standard DE, the fitness of the final trial individual *O*[*i*] is compared with the fitness of the corresponding target individual *P*[*i*], to determine which one enters the population in the next generation.

The settings used to control the RosettaDOCK local search are shown in Supplementary Information and a comparison between different settings (found in Supplementary Information) has been analyzed in terms of energy sampling (Table S4), computational time (Supplementary Table S5) and how the energy improvement at candidates achieved by the Local Search method contributes to the evolutionary process (Figure S4).

### 4.4. RosettaDOCK

Fixed-backbone RosettaDOCK models were obtained using the settings recommended in a previous benchmark study (Chaudhury et al., 2011). The method starts with randomization of the initial position by using the Rosetta parameters *randomize1*, *randomize2* and *spin*. The *nstruct* parameter, which defines the desired number of solutions to be obtained, was set to 100, 000 in order to generate the model dataset. For each solution, RosettaDOCK performs 500 MMC cycles using the coarse-grained representation and 50 MMC cycles with the full-atom representation (the local search algorithm used at EvoDOCK) but with a greedy sidechain optimization replacing the combinatorial step in 7 out of 8 cycles. The command line and further explanation parameters, used for RosettaDOCK can be found in Supplementary Information.

### 4.5. Setup for local docking

Local docking with EvoDOCK follows the same approach used with global docking. Local docking assumes a known starting binding mode and constrains the sampling search around it. In local docking with RosettaDOCK a maximum rotation perturbation of 3 and a maximum translation of 8 Å are typically used by using the flag *dock_pert 3 8*. The command line for the docking perturbation can be found in Supplementary Information. Since EvoDOCK is a population-based search, the random population created using a local perturbation approach may include a nearby solution to the “native” complex. In order to avoid the fast convergence of the algorithm to that given solution, the initial solutions are sampled with the same maximum perturbations (3 for rotation and 8 Å for translation), followed by filtering the starting points to avoid models with an RMSD to the starting model of less than 2 Å. Both methods are used to find 5,000 optimized models, EvoDOCK by running 50 independent runs with 100 individuals, while RosettaDOCK “nstruct” parameter is set to 5,000.

### 4.6. Setup for flexible backbone docking

Flexible backbone docking was carried out using a conformational ensemble with 100 members for both receptor and ligand based on the structure of the unbound proteins (PDB IDs *1kwm* and *2jto*). Following the methodology in (Marze et al., 2018), the relax, normal mode analysis and backrub protocol in Rosetta was used to create 30, 40 and 30 structures, respectively (command lines in supplementary materials). For local docking the ensemble members were superimposed onto the bound complex prior to docking. 5000 docking models were produced using these ensembles using local docking in RosettaDOCK following (Marze et al., 2018) and compared to 5000 models generated by local docking with EvoDOCK. EvoDOCK follows a similar strategy to RosettaDOCK. The initial population starts with random pairs selected from the conformational ensemble. During evolution, EvoDOCK samples the rotation and translation as in the fixed-backbone sampling by Differential Evolution and a local search. At the end of each generation, a backbone swap is performed in the population. For each individual the backbone of the receptor or ligand is swapped with a conformer from the ensemble with a given “swap probability” (10%in this study). Once the new backbone is introduced, a local search is performed based on this new starting point. Finally, if the energy is improved the new backbone replaces the old one in the individual for the next round.

### 4.7. Probabilistic measure for docking

For each target complex, we generated re-sampled decoy sets and calculated bootstrap statistical measures based on the observed docking results from each re-sampled decoy set. We generated B resampled decoy sets with replacement. From each N set of re-sampled decoy sets we calculated the probability of observing a successful docking result (lowest energy solution with iRMSD lower than 2 Å) and calculated the average for a number of resampled datasets of 5,000 (*B* = 5000) datasets as follows:

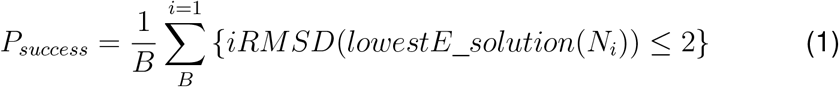

The computational time of the sampled dataset was calculated by multiplying the number of decoys/runs with the average time expended to obtain an independent trajectory with the given method (RosettaDOCK or EvoDOCK). The time to obtain an average model for EvoDOCK and RosettaDOCK is found in Supplementary Information, Table S6.

## 5. Software and availability

EvoDOCK was developed in Python 3, using PyRosetta (Chaudhury et al., 2010) version 4 and can be used a standalone program with command line. The EvoDOCK source files are available at https://github.com/Andre-lab/evodock.

