## Supplementary Information for "A memetic algorithm enables efficient local and global all-atom protein-protein docking with backbone and sidechain flexibility"

Daniel Varela, Vera Karlin and and Ingemar André
*Division of Biochemistry and Structural Biology, Department of Chemistry, Lund University, POB 124, SE-22100 Lund, Sweden*

March 11, 2022

### Supplementary Information Command lines.

#### EvoDOCK command line.

EvoDOCK runs using the following command line with a configuration file such as the example shown below. The definition of the input parameters is commented in the configuration file.

```
$ python evodock.py configuration.ini
```

```
[Docking]
# selects docking protocol [Global, Local, Unbound, RefineCluspro]
type=Global

[inputs]
# pdb, secondary structure file and fragment files
pose_input=INPUT.pdb
# bound complex used for metrics such as RMSD
native_input=BOUND_COMPLEX.pdb

# in case of Unbound docking, input list of ligand and prepacked PDBs
path_ligands=LIGAND_PREPACKED_PDBS*.pdb.ppk
path_receptors=RECEPTOR_PREPACKED_PDBS*.pdb.ppk

# in case of Cluspro Refinement, input pdb list to refine
cluspro_pdbs=INITIAL_PREPACKED_PDBS*.pdb

[outputs]
# output folder and evolution file log
output_file=/resultados/evolution_sample.log

[DE]
# evolution algorithm parent strategy [RANDOM, BEST]
scheme=RANDOM
# population size
popsize=100
# mutation rate (weight factor F)
mutate=0.9
# crossover probability (CR)
recombination=0.3
# maximum number of generations/iterations (stopping criteria)
maxiter=100
# recommended local search strategy uses simplified version of MC-based RosettaDOCK
local_search=mcm_rosetta
```

Listing 1: configuration.ini

The number of the Metropolis Monte Carlo cycles can be controlled with the Rosetta flags as follows:

#### EvoDOCK Rosetta flags for MC local search

```
# flags for local search operator
-docking:dock_mcm_first_cycles 1 /
-docking:dock_mcm_second_cycles 1
```

#### Preparation of structures

The input complex in PDB format using "-s" parameter and the static molecule and the moving molecule chains are indicated by using "-partners" as "chain1" and "chain2" respectively. Parameter "*−dock_ppk*" runs the prepack application of Rosetta, while "-ex1", "-ex2aro" and "-unboundrot" are used to include extra rotamers that would be optimized at all the residues by using "-extrachi_cutoff 0".

```
# prepack protocol command line
$ docking_protocol.linuxgccrelease /
-database $database_path /
-s $name.pdb /
-partners $chain1_$chain2 /
-dock_ppk /
-ex1 -ex2aro /
-extrachi_cutoff 0 /
-unboundrot $name.unboundrot.pdb
```

#### RosettaDOCK global docking

The input complex in PDB format using "-s" parameter and the static molecule and the moving molecule chains are indicated by using "-partners" as "chain1" and "chain2" respectively. By using "-randomize1", "-randomize2" and "-spin", both molecules are placed at random positions. On this way, initial positions for docking are randomized to perform the global docking without any input information.

```
# command line for global docking
$ docking_protocol.linuxgccrelease /
-database $database_path /
-s $name.prepack.pdb /
-nstruct $nstruct /
-partners $chains1_$chains2 /
-randomize1 /
-randomize2 /
-spin /
-ex1 -ex2aro /
-use_input_sc /
-unboundrot $name.unboundrot.pdb
```

#### RosettaDOCK local docking

The input complex in PDB format using "-s" parameter and the static molecule and the moving molecule chains are indicated by using "-partners" as "chain1" and "chain2" respectively. In the local docking problem, the initial positions are created by a random perturbation in the input complex ("-dock_pert") by using a maximum rotation and translation of 3 and 8 Å.

```
# command line for local docking
$ docking_protocol.linuxgccrelease /
-database $database_path /
-s $name.prepack.pdb /
-nstruct $nstruct /
-partners $chains1_$chains2 /
-dock_pert 3 8 /
```

```
-ex1 -ex2aro /
-use_input_sc /
-unboundrot $name.unboundrot.pdb
```

#### Ensemble generation

By using the following methods (relax, normal mode analysis and backrub), 100 conformational ensembles are generated for both receptor and ligand based on the structure of the unbound proteins (PDB IDs *1kwm* and *2jto*).

##### Relax

```
# command line for relax method
$ relax.default.linuxgccdebug -in:file:s <input PDB> -nstruct 30 -relax:thorough
```

##### Normal Mode Analysis

```
# command line for normal mode analysis method
  $ rosetta_scripts.default.linuxgccrelease \
  -in:file:s <input PDB> \
  -nstruct 40 \
  -parser:protocol nma.xml
```

where nma.xml is

```
  # nma.xml
<ROSETTASCRIPTS>
    <SCOREFXNS>
        <ScoreFunction name="bn15_cart" weights="beta_nov15_cart" />
    </SCOREFXNS>
    <RESIDUE_SELECTORS>
    </RESIDUE_SELECTORS>
    <TASKOPERATIONS>
    </TASKOPERATIONS>
    <FILTERS>
    </FILTERS>
    <MOVERS>
        <NormalModeRelax name="nma" cartesian="true" centroid="false"
scorefxn="bn15_cart" nmodes="5" mix_modes="true" pertscale="1.0"
randomselect="false" relaxmode="relax" nsample="20"
cartesian_minimize="false" />
    </MOVERS>
    <APPLY_TO_POSE>
    </APPLY_TO_POSE>
    <PROTOCOLS>
        <Add mover="nma" />
    </PROTOCOLS>
    <OUTPUT scorefxn="bn15_cart" />
</ROSETTASCRIPTS>
```

**Backrub**

```
# command line for backrub method
  $ backrub.linuxgccrelease -in:file:s <input PDB> \
  -nstruct 30 \
  -backrub:ntrials 20000 \
  -backrub:mc_kt 0.6
```

#### Prepack backbone conformers

```
# command line for docking prepack method
$ docking_prepack_protocol.linuxgccrelease \
    -in:file:s <unbound PDB >\
    -nstruct 1 \
    -ensemble1 <receptor conformer list> \
    -ensemble2 <ligand conformer list> \
    -partners A_B \
    -detect_disulf true \
    -rebuild_disulf true \
    -ex1 \
    -ex2aro
```

#### unbound-unbound docking simulations

Using the pre-packed structures, docking simulations are performed using the following command line:

```
  # command line for Rosetta unbound docking method
  $ docking_protocol.linuxgccrelease \
  -in:file:s <Pre-Packed unbound PDB> \
  -in:file:native <Bound-state PDB> \
  -nstruct 5000 \
  -ensemble1 <receptor conformer list> \
  -ensemble2 <ligand conformer list> \
  -partners A_B \
  -dock_pert 3 8 \
  -spin \
  -detect_disulf true \
  -rebuild_disulf true \
  -ex1 \
  -ex2aro
```

**Fig. S1.**

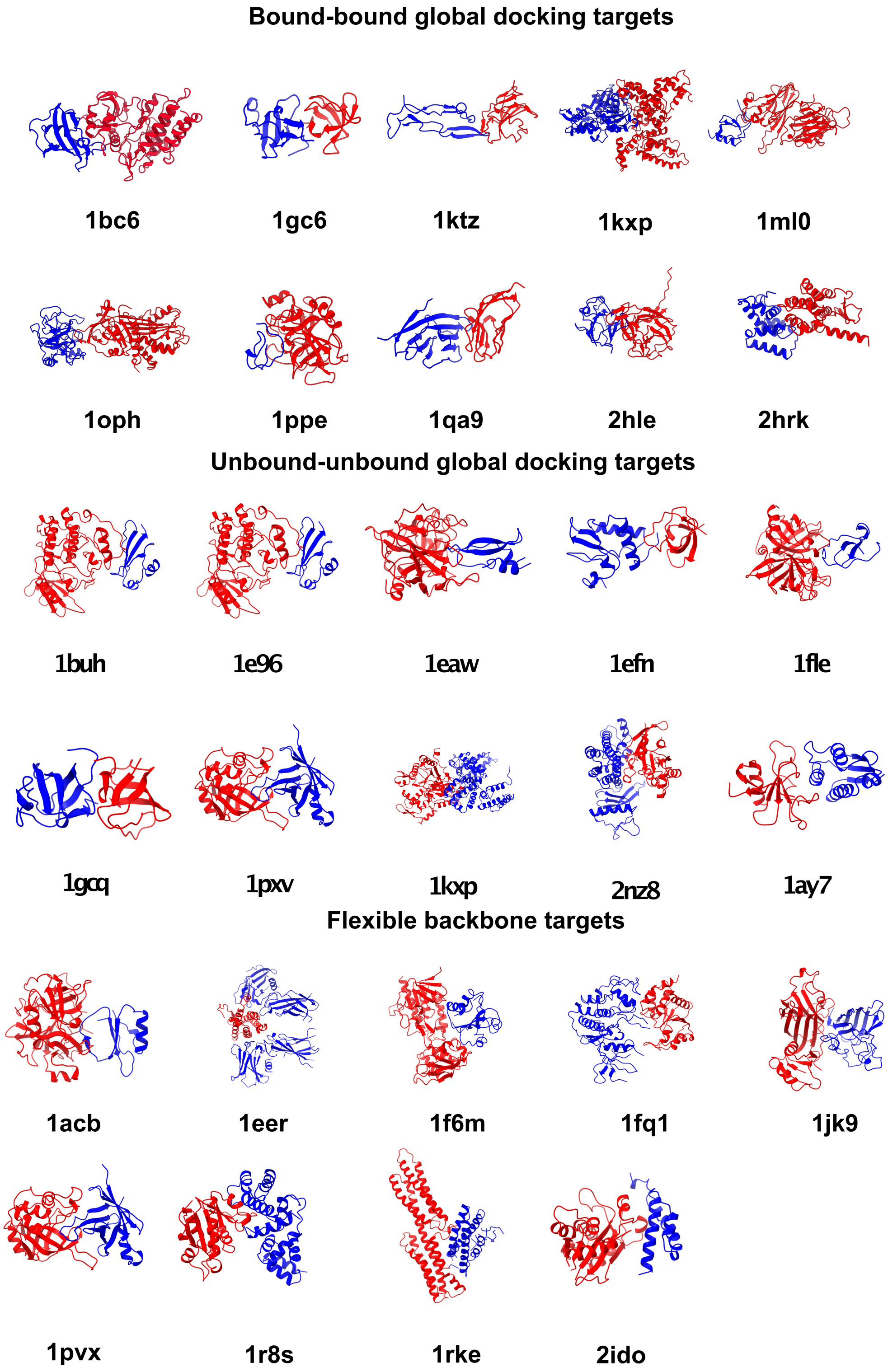


Figure 1: **Benchmark complex structures**. Complex structures used at the benchmark study. Rigid molecule is show in blue, while moving molecule is in red. Each complex is labeled by its PDB ID available at the protein data bank (RCSB).

#### Fig. S2.

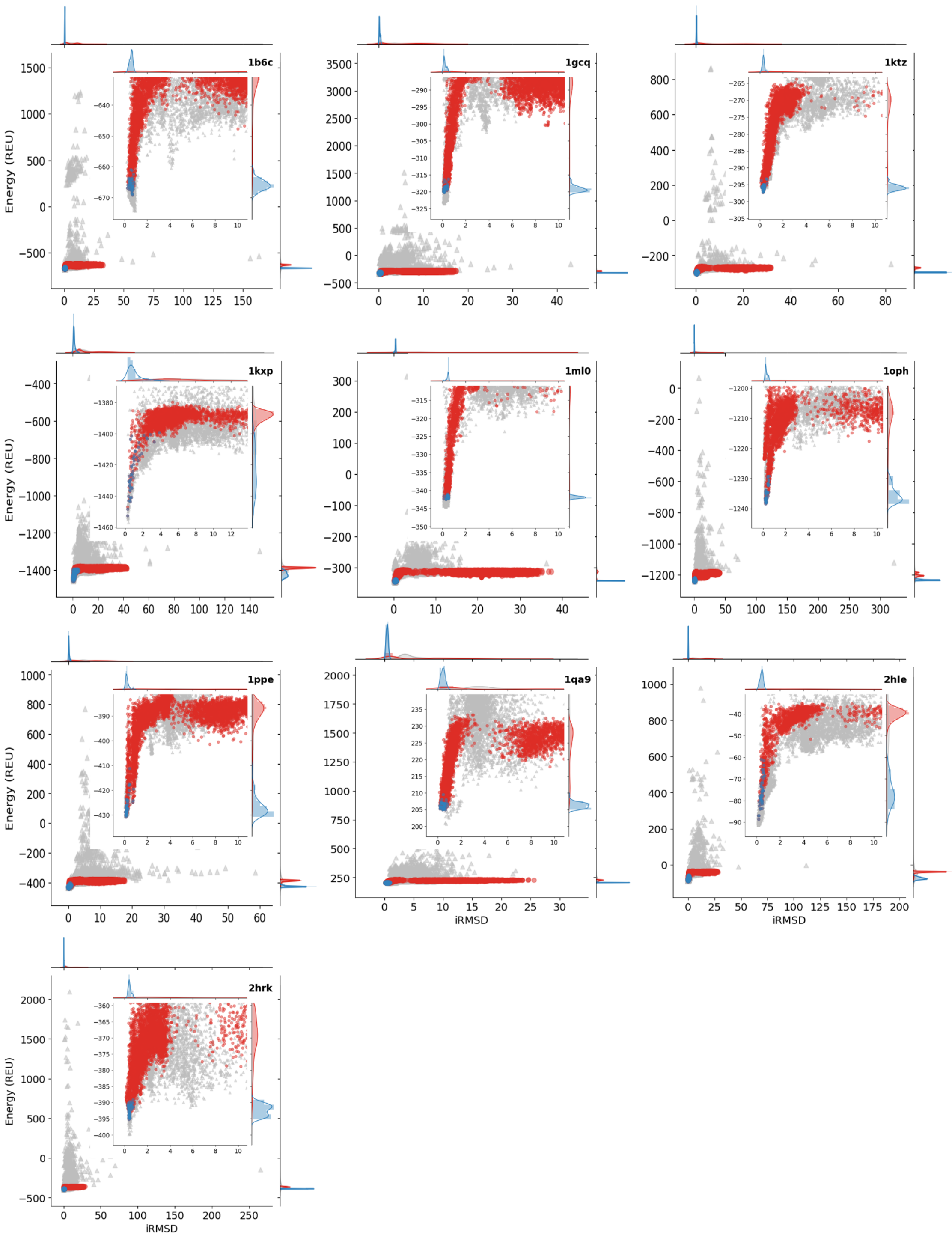


Figure 2: **Distribution of the local docking results for 5,000 models**. Each model is represented by a point where the Rosetta full-atom Energy Units (REU) are show in the $y$-axis and its interface RMSD (iRMSD) in the $x$-axis. RosettaDOCK (grey) shows the values for 5,000 independent trajectories, while EvoDOCK models (red) correspond to the optimized population of 50 independent evolutionary processes (the lowest energy solution achieved by each of the 50 runs is represented in blue). Each plot includes an additional view, zoomed into the region of low iRMSD, in order to visualize the details of "near-native" decoys.

### Fig. S3.

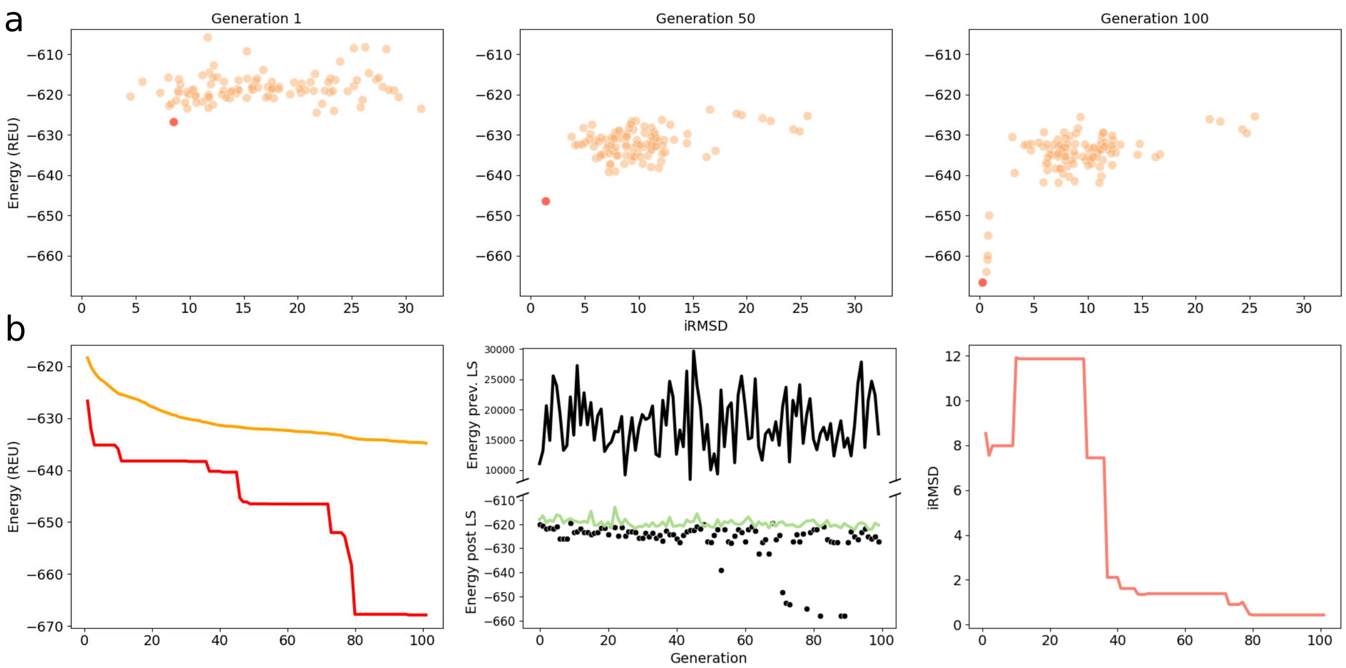


Figure 3: **Effect of Local Search across the Evolutionary Process**. **Panel a**: it shows the distribution of the Rosetta Energy in the sampled models ($y$-axis) vs. iRMSD ($x$-axis), in three snapshots in a EvoDOCK run at generations 1, 50 and 100 (left, center and right plots). Population is shown in orange plots, while the lowest energy individual is in red. **Panel b**: the left figure shows the lowest (red) and average (orange) energy values ($y$-axis) across the generations of the evolution process ($x$-axis). The figure in the center highlights the effectiveness of the Local Search operator: although DE candidates can occasionally achieve low energy values (black dots), the average energy value (black line) is very high. Therefore, the application of local search as a repair mechanism on those candidates, considerably lowers the average energy (green line), allowing the evolutionary process to obtain candidates with feasible energy for the selection procedure and lead to optimized populations. Finally, the right figure shows the evolution of the iRMSD ($y$-axis) corresponding to the solution with the lowest energy in each generation. The large changes suggested that the exploration of the method is able to avoid stagnation at a local minimum.

#### Fig. S4.

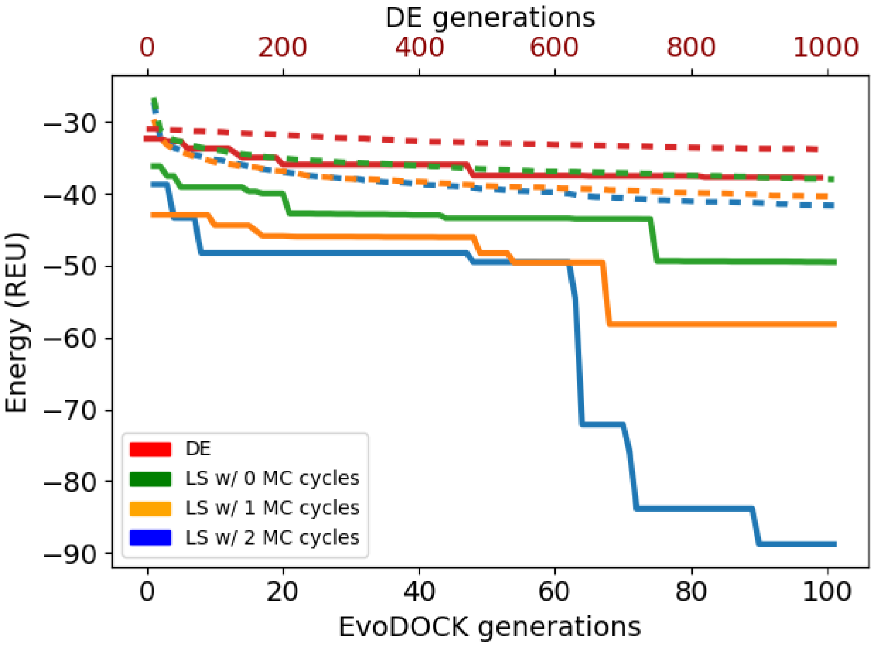


Figure 4: **Comparison of MMC cycles across the evolutionary process**. The figure shows the energy values ($y$-axis) across generations ($x$-axis), corresponding to the run which obtains the lowest energy model for complex *2hle* and by different Local Search (LS) configurations. Evolutions with 2, 1 and 0 MMC cycles are shown in blue, orange and green respectively, over 100 generations (bottom $x$-axis). Meanwhile, standard Differential Evolution without local search is shown in red over 1,000 generations (upper $x$-axis). The increase in the number of MMC cycles demostrates that the local search allows the hybrid strategy of EvoDOCK to sample lower energy values in less generations. While DE plateaus at high energy values, suggesting that the candidates created by EvoDOCK do not contribute enough to an efficient sampling of the energy landscape.

### Fig. S5.

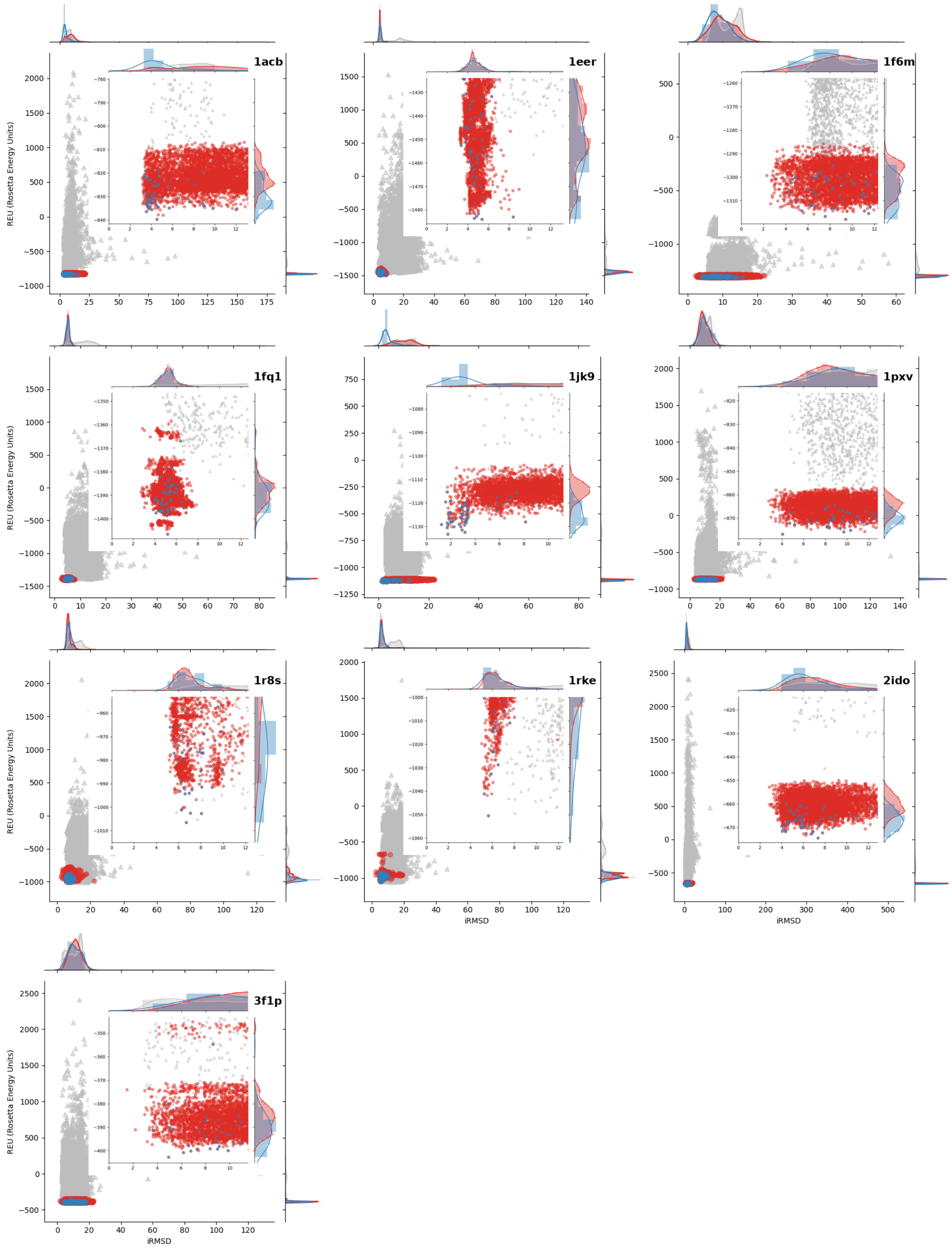


Figure 5: **Distribution of the global docking with flexible backbone results for 10,000 models**. Each model is represented by a point where the Rosetta full-atom Energy Units (REU) are show in the $y$-axis and its interface RMSD (iRMSD) in the $x$-axis. RosettaDOCK (grey) shows the values for 10,000 independent trajectories, while EvoDOCK models (red) correspond to the optimized population of 100 independent evolutionary processes (the lowest energy solution achieved by each of the 100 runs is represented in blue). Each plot includes an additional view, zoomed into the region of low iRMSD, in order to visualize the details of "near-native" decoys.

**Fig. S6.**

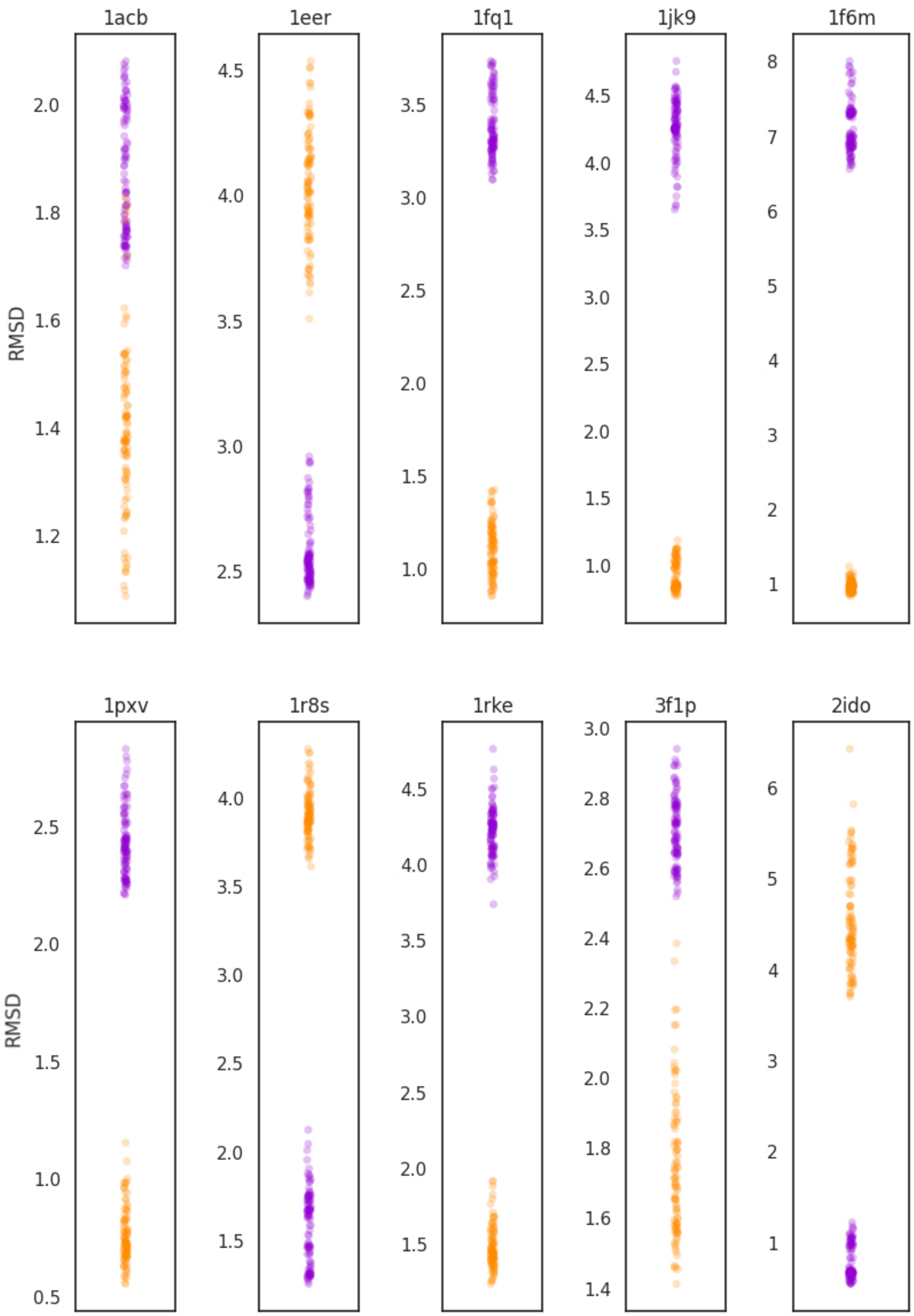


Figure 6: **Distribution of the generated ensembles**. For each protein complex, each point represents one out of 100 generated ensembles and its corresponding RMSD to the bound conformation (receptor in purple and ligand in orange).

### Table S1.

| PDB ID | EvoDOCK | RosettaDOCK |
|---|---|---|
| 1b6c | 184 (18.40%) | 7 (0.01%) |
| 1gcq | 100 (10%) | 35 (0.04%) |
| 1ktz | 220 (22%) | 25 (0.03%) |
| 1kxp | 2 (0.20%) | 0 (0%) |
| 1ml0 | 180 (18%) | 6 (0.01%) |
| 1oph | 46 (4.60%) | 3 (0%) |
| 1ppe | 227 (22.70%) | 28 (0.03%) |
| 1qa9 | 52 (5.20%) | 6 (0.01%) |
| 2hle | 3 (0.30%) | 12 (0.01%) |
| 2hrk | 197 (19.70%) | 5 (0.01%) |

Table 1: **Sampling near native models**. A near native model is defined by a threshold of 2.0 interface RMSD (iRMSD) between the sampled model and the structure deposited in the protein data bank. For each complex structure (column 1), the table shows the number of near native models and the percentage (in parentheses) of the model set obtained by independent trajectories with EvoDOCK (column 2) and RosettaDOCK (column 3).

### Table S2.

| PDB ID | $threshold = 0.6$ | $threshold = 0.8$ | $threshold = 0.95$ |
|---|---|---|---|
| 1b6c | 20.39 | 18.35 | 16.99 |
| 1gcq | 3.77 | 3.76 | 3.97 |
| 1ktz | 9.45 | 9.99 | 9.93 |
| 1kxp | $\infty$ | - | - |
| 1ml0 | 38.32 | 35.85 | 38.44 |
| 1oph | 14.68 | 14.72 | $\infty$ |
| 1ppe | 5.54 | 4.84 | 5.40 |
| 1qa9 | 33.69 | inf | $\infty$ |
| 2hle | -3.40 | -3.40 | -2.93 |
| 2hrk | $\infty$ | $\infty$ | $\infty$ |

Table 2: **Time ratio for different $P_{success}$ thresholds**. Given the amount average time neccesary to obatin a particular threshold by EvoDOCK and RosettaDOCK, the time ratio is calculated by $time(EvoDOCK)/time(RosettaDOCK)$ for each complex structure by setting a $P_{success}$ at 0.6, 0.8 and 0.95 at columns 2 to 4.

### Table S3.

| PDB ID | EvoDOCK | RosettaDOCK |
|---|---|---|
| 1b6c | -669 (0.7) | -674 (0.9) |
| 1gcq | -320 (0.2) | -324 (0.2) |
| 1ktz | -297 (0.3) | -296 (0.3) |
| 1kxp | -1453 (0.3) | -1457 (0.4) |
| 1ml0 | -342 (0.3) | -345 (0.3) |
| 1oph | -1238 (0.2) | -1237 (0.2) |
| 1ppe | -431 (0.1) | -431 (0.1) |
| 1qa9 | 205 (0.4) | 201 (0.7) |
| 2hle | -89 (0.2) | -91 (0.2) |
| 2hrk | -395 (0.4) | -400 (0.4) |

Table 3: **Local Docking energy summary**. Comparison of the lowest energy model and the corresponding and corresponding iRMSD value (in parentheses) obtained with EvoDOCK and RosettaDOCK.

### Table S4.

| PDB ID | Local Search 2 MMC cycles | Local Search 1 MMC cycles | Local Search 0 MMC cycles | Non Local search |
|---|---|---|---|---|
| 1b6c | -669 (0.6) | -670 (0.6) | -668 (0.6) | -628 (24.1) |
| 1gcq | -320 (0.1) | -319 (0.2) | -320 (0.4) | -289 (10.1) |
| 1ktz | -297 (0.3) | -297 (0.3) | -293 (0.5) | -274 (8.7) |
| 1kxp | -1413 (18.6) | -1413 (2.5) | -1413 (18.6) | -1388 (16.7) |
| 1ml0 | -343 (0.3) | -343 (0.3) | -342 (0.3) | -313 (15.2) |
| 1oph | -1238 (0.2) | -1236 (0.2) | -1222 (3.6) | -1190 (16.5) |
| 1ppe | -431 (0.1) | -431 (0.1) | -431 (0.1) | -392 (8.4) |
| 1qa9 | 205 (0.5) | 205 (0.5) | 217 (1.1) | 226 (14.0) |
| 2hle | -74 (0.3) | -58 (1.4) | -50 (8.8) | -41 (12.2) |
| 2hrk | -395 (0.4) | -395 (0.4) | -386 (0.1) | -364 (13.8) |

Table 4: **Energy sampling results with different Local Search (LS) and MMC cycles**. The lowest energy value (and the corresponding iRMSD of the model in parentheses) for different Local Search (LS) configurations are shown: using 2, 1 and 0 Metropolis Monte Carlo (MMC) energy minimization cycles (columns 2 to 4) and by using the default Differential Evolution (DE) algorithm without any Local Search method (column 5). That is, in this last case, it does not apply any repair mechanism to the DE candidates, such as *slide_into_contact* or side-chain optimization. In order to a fair comparison in terms of computational cost, The number of DE generations are increased until a similar computational time is reached (that corresponding to the 100 generations with EvoDOCK).

### Table S5.

| PDB ID | Local Search 2 MMC cycles | Local Search 1 MMC cycles | Local Search 0 MMC cycles | Non Local search |
|---|---|---|---|---|
| 1b6c | 17743 | 20013 | 12593 | 2292 |
| 1gcq | 7011 | 13952 | 3712 | 559 |
| 1ktz | 7297 | 6809 | 4891 | 823 |
| 1kxp | 35310 | 8910 | 20894 | 3588 |
| 1ml0 | 15291 | 16378 | 10727 | 2169 |
| 1oph | 27049 | 10939 | 15453 | 3159 |
| 1ppe | 11882 | 26531 | 6630 | 1225 |
| 1qa9 | 7888 | 10667 | 5320 | 920 |
| 2hle | 14215 | 7266 | 8093 | 1536 |
| 2hrk | 11840 | 5199 | 8243 | 1447 |

Table 5: **Computational time (seconds) with different Local Search (LS) and MMC cycles**. Average computational time for each structure complex with different Local Search configurations by using 2, 1 and 0 Metropolis Monte Carlo (MMC) cycles (shown in columns 2 to 4, respectively) and with the standard Differential Evolution (DE) algorithm. Average time for all benchmark complexes with the Memetic Algorithm (hybridization of DE and the MMC local search) is 4.3, 3.5 and 2.6 hours with 2, 1 and 0 minimization cycles, respectively. Although the average time is decreased as the cycles are reduced, that decrease is not uniform across all the benchmark complexes, suggesting that the minimization of cycles also contributes to the convergence of the evolutionary algorithm. DE has an average time of 30 minutes, which shows that the repair mechanism used in the local search is significantly costly, but with a great deterioration when obtaining low energies, as it can be seen in the Supplementary Table 4.

### Table S6.

| PDB ID | EvoDOCK | RosettaDOCK |
|---|---:|---:|
| 1b6c | 17743 | 143 |
| 1gcq | 7011 | 37 |
| 1ktz | 7297 | 50 |
| 1kxp | 35310 | 271 |
| 1ml0 | 15291 | 153 |
| 1oph | 27049 | 210 |
| 1ppe | 11882 | 68 |
| 1qa9 | 7888 | 62 |
| 2hle | 14215 | 113 |
| 2hrk | 11840 | 88 |

Table 6: Average computational time (in seconds) for each benchmark complex (column 1) with EvoDOCK (column 2) and RosettaDOCK (column 3).

#### Table S7.

| PDB ID | time (hours) | balanced | elec. | hydro. | vdw. |
|---|---|---|---|---|---|
| 1b6c | 8:41:40 | 1.8 (1.8) | 3.9 (1.7) | 1.9 (1.9) | 20.0 (13.5) |
| 1gcq | 2:53:00 | 10.5 (1.1) | 10.4 (1.2) | 11.2 (1.1) | 13.8 (4.0) |
| 1ktz | 5:47:40 | 20.5 (10.5) | 14.5 (1.2) | 15.0 (10.4) | 12.0 (1.6) |
| 1kxp | 23:52:00 | 1.3 (1.3) | 1.5 (1.5) | 2.0 (2.0) | 2.4 (2.4) |
| 1ml0 | 9:09:40 | 23.1 (1.0) | 27.8 (20.9) | 22.3 (1.1) | 31.2 (27.8) |
| 1oph | 11:23:20 | 1.6 (1.6) | 1.6 (1.6) | 11.1 (1.6) | 20.5 (19.7) |
| 1ppe | 4:34:40 | 1.3 (1.3) | 1.3 (1.3) | 1.2 (1.2) | 14.3 (1.2) |
| 1qa9 | 5:01:00 | 21.1 (17.5) | 21.3 (1.4) | 19.7 (17.3) | 1.5 (1.5) |
| 2hle | 7:53:40 | 2.1 (2.1) | 1.8 (1.8) | 7.5 (2.0) | 1.4 (1.4) |
| 2hrk | 7:48:20 | 2.6 (2.6) | 3.0 (3.0) | 3.6 (2.4) | 21.6 (14.9) |

Table 7: **Data results with ClusPro server**. Column 2 shows the average time in hours for PIPER with each protein complex in Column 1 (using the 'Balanced' energy scheme). From column 3 to 6, table shows the iRMSD (Å) of the representative model against the native conformation and, in parentheses, the minimum iRMSD (Å) value of all the representative models from clusters for each of the four score schemes: 'Balanced', 'Electrostatic-favored' (elec.),'Hydrophobic-favored' (hydro.) and 'VdW+Elec' (vdw)..

#### Table S8.

| PDB id | EvoDOCK | | ClusPro | |
|---|---|---|---|---|
| | DockQ | iRMSD | DockQ | iRMSD |
| 1ay7 | 0.87 (0.87) | 0.45 (0.45) | 0.69 (0.03) | 1.63 (11.83) |
| 1buh | 0.07 (0.03) | 7.21 (11.89) | 0.15 (0.03) | 4.76 (12.31) |
| 1E96 | 0.31 (0.04) | 3.5 (7.98) | 0.15 (0.14) | 4.68 (6.5) |
| 1eaw | 0.11 (0.02) | 6.68 (12.27) | 0.34 (0.07) | 2.75 (8.29) |
| 1efn | 0.36 (0.03) | 2.6 (12.36) | 0.1 (0.05) | 5.63 (11.85) |
| 1pxv | 0.08 (0.02) | 7.3 (13.42) | 0.56 (0.02) | 1.79 (13.74) |
| 2nz8 | 0.1 (0.03) | 8.1 (15.56) | 0.05 (0.01) | 10.41 (24.61) |
| 3hi6 | 0.1 (0.03) | 6.35 (11.46) | 0.22 (0.04) | 5.04 (7.07) |
| 4m3k | 0.12 (0.02) | 5.85 (14.33) | 0.05 (0.01) | 10.72 (18.29) |
| 1fle | 0.24 (0.01) | 4.08 (19.66) | 0.54 (0.28) | 1.94 (3.24) |

Table 8: **Data results of EvoDOCK and ClusPro Server at unbound docking experiments**. For each benchmark complex (Column 1), columns 2-3 shows the results of EvoDOCK for DockQ and iRMSD, with best value of the lowest energy individual for 100 different trajectories (while in the parentheses it is shown the value of the lowest energy model). Columns 4 to 6 shows the DockQ and iRMSD values for the ClusPro server, for all the obtained models and, in parentheses, the model from the most representive cluster using 'Balanced' energy scheme, as it is done at CAPRI experiments.

#### Table S9.

| PDB id | EvoDOCK | | | RosettaDOCK | | |
|---|---|---|---|---|---|---|
| | DockQ | iRMSD | CAPRI | DockQ | iRMSD | CAPRI |
| 3f1p | 0.52 | 2.62 | Medium | 0.48 | 2.61 | Acceptable |
| 1r8s | 0.18 | 5.63 | Incorrect | 0.05 | 7.93 | Incorrect |
| 1eer | 0.45 | 2.53 | Acceptable | 0.33 | 3.37 | Acceptable |
| 2ido | 0.27 | 4.76 | Acceptable | 0.39 | 3.63 | Acceptable |
| 1pxv | 0.21 | 4.67 | Incorrect | 0.36 | 3.69 | Acceptable |
| 1fq1 | 0.47 | 3.31 | Acceptable | 0.43 | 3.54 | Acceptable |
| 1acb | 0.46 | 2.48 | Acceptable | 0.40 | 2.98 | Acceptable |
| 1jk9 | 0.81 | 0.99 | High | 0.54 | 3.05 | Medium |
| 1rke | 0.35 | 5.36 | Acceptable | 0.31 | 5.51 | Acceptable |
| 1f6m | 0.28 | 5.81 | Acceptable | 0.24 | 6.12 | Acceptable |

Table 9: **Data results of EvoDOCK and RosettaDOCK at unbound flexible backbone docking**. For each benchmark complex (Column 1), columns 2-4 shows the results of DockQ, iRMSD and CAPRI classification for the lowest energy models of 100 different trajectories with EvoDOCK. Columns 5 to 7 shows the results of the 100 lowest energy models sampled at 10,000 trajectories of RosettaDOCK.
